# Comparative dissection of the peripheral olfactory system of the Chagas disease vectors *Rhodnius prolixus* and *Rhodnius brethesi*

**DOI:** 10.1101/2021.01.08.425861

**Authors:** Florencia Campetella, Rickard Ignell, Rolf Beutel, Bill S. Hansson, Silke Sachse

**Affiliations:** Department of Evolutionary Neuroethology, Max Planck Institute for Chemical Ecology, Hans-Knoell-Str. 8, 07745 Jena, Germany; Unit of Chemical Ecology, Department of Plant Protection Biology, Swedish University of Agricultural Sciences, SE-23053 Alnarp, Sweden; Institute for Zoology and Evolutionary Biology, Friedrich Schiller University, Jena, Germany

## Abstract

American trypanosomiasis or Chagas disease is thought to be transmitted by both domestic and sylvatic species of Triatominae. These haematophagous insects use sensory cues to find their vertebrate hosts. Among them, odorants have been shown to play a key role. Previous work revealed morphological differences in the sensory apparatus of sylvatic and domestic species of Triatomines, but to date a functional study of the olfactory system is not available. After examining the antennal sensilla with scanning electronic microscopy (SEM), we compared olfactory responses of the domestic *Rhodnius prolixus* and the sylvatic *Rhodnius brethesi* with an electrophysiological approach. In electroantennogram (EAG) recordings, we first show that the antenna of *R. prolixus* shows high responses to carboxylic acids, compounds found in their habitat and headspace of hosts. We then compared responses from olfactory sensory neurons (OSNs) housed in the grooved peg sensilla of both species as these are tuned to these compounds using single-sensillum recordings (SSR). In *R. prolixus*, the SSR responses revealed a narrower tuning breath than its sylvatic counterpart, with the latter showing responses to a broader range of chemical classes. Additionally, we observed significant differences between these two species in their response to particular volatiles, such as amyl acetate and butyryl chloride. In summary, the closely related, but ecologically differentiated *R. prolixus* and *R. brethesi* display distinct differences in their olfactory functions. Considering the ongoing rapid destruction of the natural habitat of sylvatic species and likely shifts towards environments shaped by humans, we expect that our results will contribute to the design of efficient vector control strategies in the future.

**Author Summary:** American Tripanosomiasis, also known as Chagas disease, is a disease which no one speaks out, although there are up to eight million people infected worldwide. Its causative agent is the protozoan *Tripanosoma cruzi* which is transmitted by triatomine insects, alias kissing bugs. Several studies have highlighted the importance of olfaction for host-seeking behavior in these insects, which enables them to target their vertebrate hosts, and to get their vital blood meal, while infecting them at the same time. Vector control strategies have been the most efficient policy to combat the spread of Chagas disease by triatomine insects. However, recent changes in the natural habitats of these insects challenge their effectiveness, as species so far thought to be exclusive to sylvatic environments are now frequently found in peridomestic areas. In this context, to understand how sylvatic and domestic kissing bugs detect odors to locate their host and choose their habitats is highly relevant. In this study, we compare the olfactory system of the domestic kissing bug *Rhodnius prolixus* and its sylvatic counterpart *Rhodnius brethesi* at a morphological and functional level. We reveal that detection of host and habitat volatiles share many similarities, but also exhibit pronounced differences between both species.

## Introduction

Chagas or American trypanosomiasis, caused by infection with the protozoan *Trypanosoma cruzi*, is a chronic disease that is endemic in 21 Latin American countries, where it significantly affects the most vulnerable inhabitants. It is estimated that its prevalence in some areas can be as high as 5%, and its annual burden in health care costs sums up to 600 million dollars [1]. Already in 1905 it was shown that blood-sucking insects belonging to the Triatominae subfamily (Heteroptera: Reduviidae) transmit *T. cruzi* through their faeces. To date, the most effective and successful methods to control the spread of American trypanosomiasis have been vector control policies. Wide-spread use of pesticides and training of local communities to identify and kill the insects are the most efficient strategies to date [2]. However, with the appearance of pesticide-resistant insects, new management strategies are urgently needed.

Triatominae is a poorly defined and possibly paraphyletic group of the predaceous true bugs of the family Reduviidae [3]. All 151 described species, phylogenetically grouped into five tribes [4], are capable of transmitting Chagas disease [5]. From these, some species, such as *Rhodnius prolixus* and *Triatoma infestans*, are considered particularly important from an epidemiological standpoint, as they are often found associated with households (*i.e*., domestic species). However, most of the species of the Rhodniini tribe, to which *R. prolixus* belongs, are sylvatic, and found in the forest, nesting on palm trees [6]. While some species are reported to nest in a limited number of specific palm tree species, and are thus thought to be specialists, others, the generalists, find shelter in more than one palm species [7–9]. *R. prolixus* is known to be of this latter type. An interesting example of a specialist species is *Rhodnius brethesi*, which, so far, has only been found on the palm tree species *Leopoldina piassaba* [8, 9]. Despite the interesting nature of these associations, studies on sylvatic species has been marginal, with most of the research focused on domiciliated species. However, as deforestation and climate change increase [7, 10], sylvatic species will lose their natural habitats and might find refuge in domestic and peridomestic areas [11], putting its inhabitants at higher risk and becoming a public health problem. Thus, in order to design better vector control strategies, a thorough understanding of the differences and similarities between domestic and sylvatic species is needed.

Being active at night, triatomines make use of physical and chemical cues to find their hosts [12, 13]. Several studies have highlighted the importance of olfaction for host-seeking behavior in these insects [14–18]. Terrestrial vertebrates, the main host for these obligatory haematophogaous insects, emit odor signatures that can be composed of up to 1000 different volatiles [19–21], many of them being produced by the skin microflora [22]. Previous work has shown that the domestic species, *T. infestans* and *R. prolixus*, make use of some of these volatiles to find their hosts [15, 16, 18, 23–27]. In particular, carbon dioxide, 1-octen-3-ol, acetone, several amines, as well as carboxylic acids are attractive cues for *R. prolixus* [16, 28, 29]. However, how *R. prolixus* and other species detects these cues remains largely unknown.

The olfactory system of sylvatic and domestic species has been proposed to be tuned to different odor compounds due to their different ecological niches [8]. While domestic species are exposed to a limited number of volatiles in their environments, sylvatic insects need to be able to discriminate among a larger number in order to find hosts, nest and oviposition sites. Morphological studies in Triatominae have shown that the number of olfactory sensilla is correlated with the complexity and number of ecotypes in which the insects are found [30, 31]. For instance, domestic species with stable environments have lower number of chemosensory sensilla than their sylvatic relatives [30–34]. Furthermore, a reduced expression of olfactory binding proteins (OBPs) and chemosensory proteins (CSPs) are found in domiciliated insects of *Triatoma brasiliensis*, compared to sylvatic and peridomestic ones [35].

In the present study, we used a comparative approach to characterize the peripheral olfactory system of domestic and sylvatic species of triatomines at an anatomical and functional level. We selected the generalist *R. prolixus* as a domestic species, while the closely related sylvatic species *R. brethesi* represents a specialist. We investigated whether habitat choice is reflected in the olfactory system of these insects. This is, to our knowledge, the first time a functional comparative study between domestic and sylvatic triatomine species is carried out.

## Material and methods

### Insect rearing

Insects were reared as described in detail in previous publications [36, 37]. Adult males of *R. brethesi* and *R. prolixus*, starved for 3-4 weeks, were used in the experiments. Batches of insects were kept in individual boxes with a Light:Dark cycle set to 12:12h. The boxes were placed inside a chamber at 25°C and 60% relative humidity. Each insect was used at the beginning of the scotophase, as it has been shown that olfactory acuity is higher at this timepoint [15, 38].

Laboratory rearing has been shown to have a species-specific impact on the number and distribution of olfactory and mechanosensory sensilla [34]. However, according to previous work, in the case of *R. prolixus* this effect is either non-existant or only moderate [39]. In *R. brethesi* an increase in the density of mechanosensory sensilla (bristles), and a reduction in the number of trichoid and basiconic sensilla has been observed in laboratory-rerared insects compared to wild ones [33]. Despite our efforts, it was not possible to include specimens of this species from the field.

### SEM

The heads of the insects, including the antennae were fixed with 2.5% (v/v) glutaraldehyde in cacodylate buffer (pH 7.4) for 60 min. Afterwards the samples were washed three times for 10 min with cacodylate buffer and dehydrated in ascending ethanol concentrations (30%, 50%, 70%, 90% and 100%) for 10 min each. Subsequently, the samples were dried at the critical-point using liquid carbon dioxide, and sputter coated with gold (approximately 2 nm) using a SCD005 sputter coater (BAL-TEC, Balzers, Liechtenstein). Finally, the relevant surfaces were analyzed with a scanning electron microscope (SEM) LEO-1450 (Carl Zeiss NTS GmbH, Oberkochen, Germany) providing a rotating sample stage to allow all-around imaging.

### Odors

Odors were obtained from Sigma-Aldrich, FLUKA, Aldrich at the highest purity available. Compounds used are listed in Supplementary Table 1 and Supplementary Table 2, together with the respective solvent used (paraffin oil, CAS: 8012-95-1, Supelco, USA; distilled water; or ethanol, Sigma Aldrich, Germany), in which each odor was diluted. For electroantennogram (EAG) recordings, a dilution of 10% in paraffin oil (Supelco, USA) was used, while we applied all odors at a dilution of 1% in single sensillum recordings (SSRs). An odor blend, used only in SSR, was created by mixing all compounds listed in **S2 Table** in a 1:1 ratio, all at 1% dilution in paraffin oil. The compounds in this blend are known to be detected by odorant receptors (ORs) in other species, and were thus designed to identify possible ORs housed in the grooved peg sensilla of *Rhodnius spp* [40–43].

### Odor application

Odors used as stimuli were prepared at the beginning of each experimental session: a 10 μl aliquot of the diluted odor (see Supp. Table 1, Supp. Table 2) was pipetted onto a fresh filter paper (Ø=1 cm^2^, Whatman, Dassel, Germany), which was placed inside a glass Pasteur pipette. Each loaded filter paper was used for a maximum of 3 times to ensure a stable concentration across experiments. Highly volatile carboxylic acids and aldehydes were loaded at each stimulus presentation. A stimulus controller (Stimulus Controller CS-550.5, Syntech, Germany) was used to deliver odors to the insect antenna through a metal pipette placed less than 1.5 cm (EAG) or 0.5 cm (SSR) away from the insect antenna. A constant humidified air flow of 1.0 l min^−1^ was delivered to the insect, while each odor pulse had an airflow of 0.5 l min^−1^, and was buffered with compensatory airflow of the same magnitude.

### Electroantennogram (EAG) recordings

An antenna was severed quickly between the scape and the pedicel and placed between two metal electrodes. Conductive gel (Spectra 360, Parker Laboratories, Fairfield, USA) was applied to each end of the antenna. The electrode was connected to a Syntech IDAC analog/digital converter (Syntech). Acquisition was done with Autospike32 at a sample rate of 2400 Hz. While the application of odors was randomized we did ensure to apply the control (paraffin oil) at regular intervals. During the screening of the odor panel, we observed an increase in the response amplitude to the control in function of time. To account for this bias, we normalized each recording with the following formula, similarly to how it has been previously done [44]:

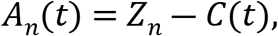

with,

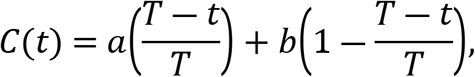

where *A_n_*(*t*) is the normalized response to a given odor stimulus *n* at a given time *t*; *Z_n_* is the measured response to the odor stimulus *n*, and *C*(*t*) is the averaged solvent response at a given time *t*, with *a* being the closest solvent response before stimulus presentation at *t_a_*, and *b* the closest solvent response after stimulus presentation at *t_b_*. The contribution of each of these solvent responses to the averaged solvent response is pondered by the factor 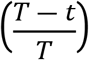, where *T* = *b − t_a_*.

### Single-sensillum recordings (SSR)

Insects were placed inside a severed 5 ml plastic tip (Eppendorf, Hamburg, Germany), which was sealed with dental wax (Erkodent, Pfalzgrafenweiler, Germany). The tip was then immobilized on a microscopy slide with dental wax. Both antennae were glued to a coverslip with double-sided tape. A tungsten electrode inserted into the insect’s abdomen was used as reference. Preliminary recordings with a silver wire as a reference electrode did not show an improvement in the signal-to-noise ratio. The preparation was placed under an upright microscope (BX51WI, Olympus, Hamburg, Germany) equipped with a 50x air objective (LMPlanFI 50x/ 0.5, Olympus). Neural activity in the form of spike trains originating from OSNs was recorded with a sharpened tungsten electrode targeted at the base of a grooved peg sensillum. Signals were amplified (Syntech Universal AC/DC Probe; Syntech) and sampled at 10,666.7 samples/s through an USB-IDAC (Syntech) connected to a computer. Spikes were extracted using Autospike32 software (Syntech). Odor responses from each sensillum were calculated as the difference in the number of impulses 0.5 s before and after stimulus onset.

The response to each odorant in the SSR recordings was calculated as the change in spikes/s upon odor stimulation using Autospike32. The response to the solvent was subtracted from each measurement. The number of OSNs housed in each sensillum in *R. prolixus* has been estimated to be between 5 and 6 [23, 45]. We attempted to confirm this observation using semi-thin section of the antenna (data not shown) but, despite our efforts, were unable to decisively identify the number of sensory neurons in the grooved peg (GP) sensilla in any of the two species. For that reason, we decided to define each sensillum as a responding unit, as it has been done in other insects [46].

Subsequent analysis was carried out in MATLAB (The MathWorks Inc, Natick, USA) in which an agglomerative hierarchical clustering of the sensillum responses, with a Euclidean metric and Ward’s method, was performed. The inconsistency coefficient was calculated for each link in the dendrogram, as a way to determine naturally occurring clusters in the data [47, 48]. A depth of 4 and a coefficient cutoff of 1.8 for *R. prolixus* and 1.0 for *R. brethesi* were used in the calculation.

In **Figure 4B** the response of each sensillum type was taken as the average response of individual sensilla belonging to the same cluster (i.e., sensillum type). In **Figure 4C** the average responses of each sensillum type were then grouped and averaged for the chemical classes. These responses were then normalized to the maximum response within each sensillum type. Principal component analysis (PCA, with a Singular Value Decomposition (SVD) algorithm) was performed in MATLAB (The MathWorks Inc) using the averaged scaled (between 0 and 1) and z-score normalized responses for each sensillum type and each species. As a measure for similarity, we applied a One-Way ANOSIM to calculate whether different sensillum types represent significantly different classes [49].

Averaged responses were computed as the mean of all sensillum responses to a particular odorant. To compare among chemical classes, these odor responses were then further averaged for each particular chemical class. Comparison between species was done using unpaired two-tailed Student t-tests (GraphPad Prism 8, San Diego, USA). These average responses were then normalized to the odor that elicited the hightest responses in each species, being for both species propionic acid, and the lifetime sparseness (S) of each sensillum was calculated. We applied the lifetime sparseness as a measure of the response breadth of each sensillum. For its calculation the following formula was used [50]:

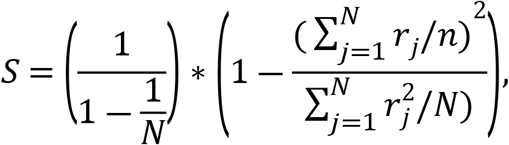

where S is the lifetime sparseness, *N* is the number of tested odors and *r_j_* is the sensillum response to any given odor *j*, with *r_j_* ≥ 0 and *S* ∈ [0,1], where *S*=0 corresponds to the case in which the sensillum responds equally to all odorants, and *S*=1 to where the sensillum responds to only one odor of the set.

## Results

### Species-specific morphological differences of the antenna

Several studies have put forward the hypothesis that sensillum patterns in haematophagous insects, including different Triatominae species, reflect specific adaptations to different hosts and habitats [51–53]. Until now, comparative qualitiative and quantitative studies of the main olfactory organ, the antenna, of the domestic species *R. prolixus* and the sylvatic specialist *R. brethesi* are lacking. We therefore aimed to investigate as a first step potential morphological differences between the antennae of both species using scanning electronic microscopy (SEM) (**Fig. 1, S1 Fig.** and **S2 Fig.**). A qualitative analysis of the morphological patterns of sensilla on the antenna of both species did not reveal major differences. In both species, the second segment, or pedicel, is enriched in sensilla described to have a mechanosensory function [38–40]: sensilla trichobothrium, bristles I and III (**S1 Fig.** and **S2 Fig.**). Additionally, a sensillum shown to have a thermo-receptive function, the cave organ, was known in *R. prolixus* and *T. infestans* [54, 55], is also present in *R. brethesi* (**S2B Fig.**). However, we were unable to identify the previously described ornamented pore [38] in the sylvatic species, possibly due to the angle of orientation of our preparation. Notably, our micrographs show the existence of two previously undescribed sensillum types on the pedicel of *R. prolixus*. One is a peg-in-pit sensillum with an inflexible socket, with no evident pores, housed within a chamber in the antennal cuticle (**S1B Fig.**). This type is reminiscent of the thermosensitive sensilla coeloconica [42], but its function remains unknown. The second sensillum type resembles a type 3 coeloconic sensilla, characterized by two pores at its base, described in other hemipteran species (**S2E Fig.**) [45].

**Figure 1.**
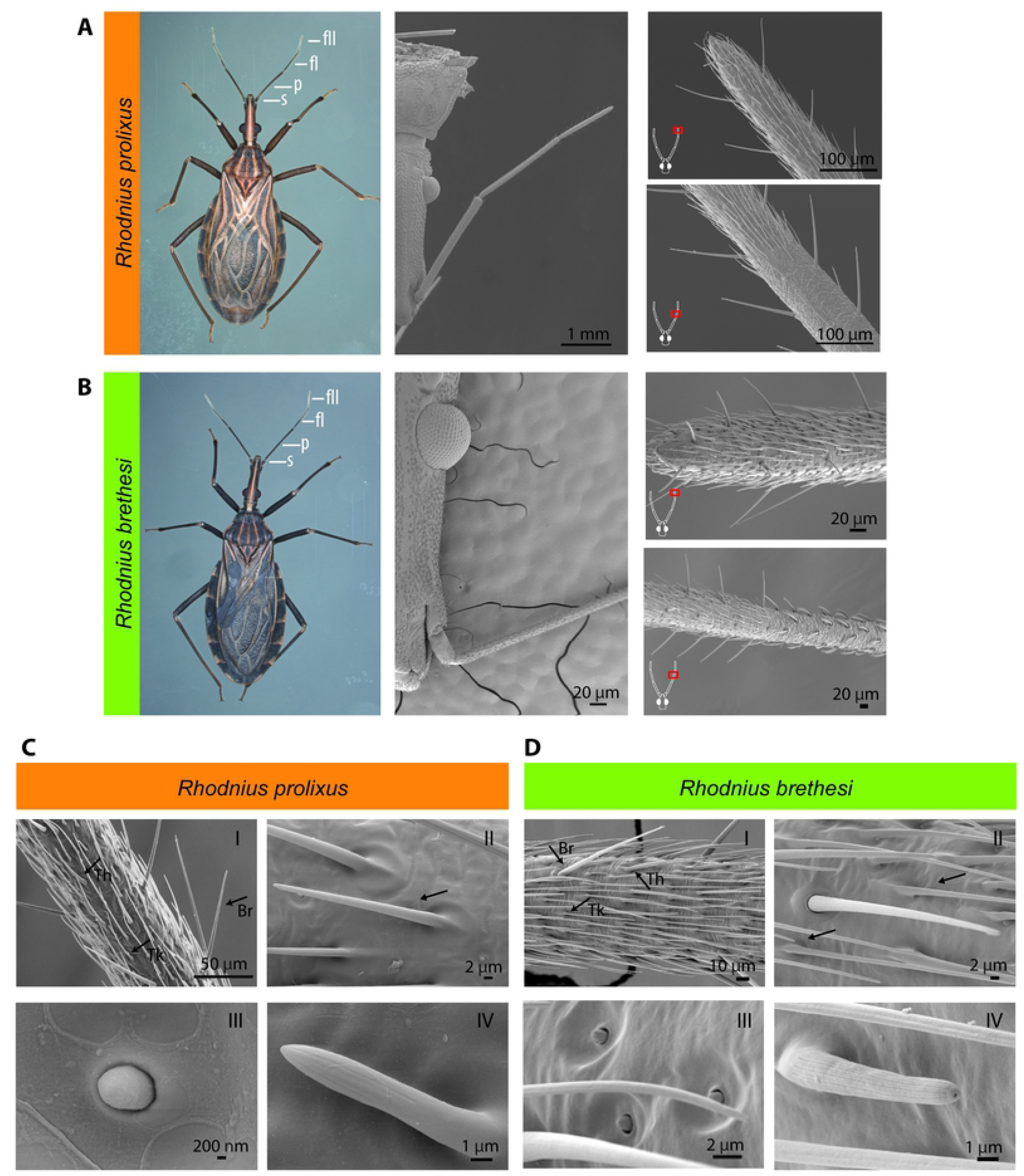
Morphology of the peripheral olfactory system of *Rhodnius prolixus* and *Rhodnius brethesi*. (A,B) Light-microscopic images of whole body of *R. prolixus* and *R. brethesi* (left panel). Antennal segments indicated: scape (s), pedicel (p), flagellomere I (fI), flagellomere II (fII). Scanning electron microscopy (SEM) images of head (middle panel) and corresponding antennal segments (right panel). (C,D) Antennal sensilla of *R. prolixus* and *R. brethesi* on flagellomere II: arrows indicate (I) thick-walled (Tk), and thin-walled (Th) trichoid sensilla, and bristles (Br). (II) Thick-walled trichoid sensillum, (III) campaniform sensillum, and (IV) grooved peg sensillum.

Wall-pore sensilla with inflexible sockets were found on flagellomere I and the distal half of flagellomere II in both species. These include the thick- and thin-walled trichoid, as well as the double-walled grooved peg sensilla (**Fig. 1C, D**). All of these sensillum types show slight longitudinal grooves filled with pores or slits indicative of an olfactory function [24]. In the same segments of both species we also identifiied a poreless sensillum with an inflexible socket: the campaniform sensillum [56, 57] (**Fig. 1C, D**).

Quantitative differences in the number of olfactory sensilla of the species potentially reflect particular adaptations to their ecological niches. Thus, we quantified the sensillum density on the flagellomere II, where putative olfactory sensilla reach the highest density and inter-specific differences were reported among triatomines [30] (**Fig. 2**). Our results show that the density of both thin- and thick-walled trichoid sensilla is significantly higher in the sylvatic *R. brethesi* than in *R. prolixus*. In contrast, the density of grooved peg sensilla is not significantly different between the two species.

**Figure 2.**
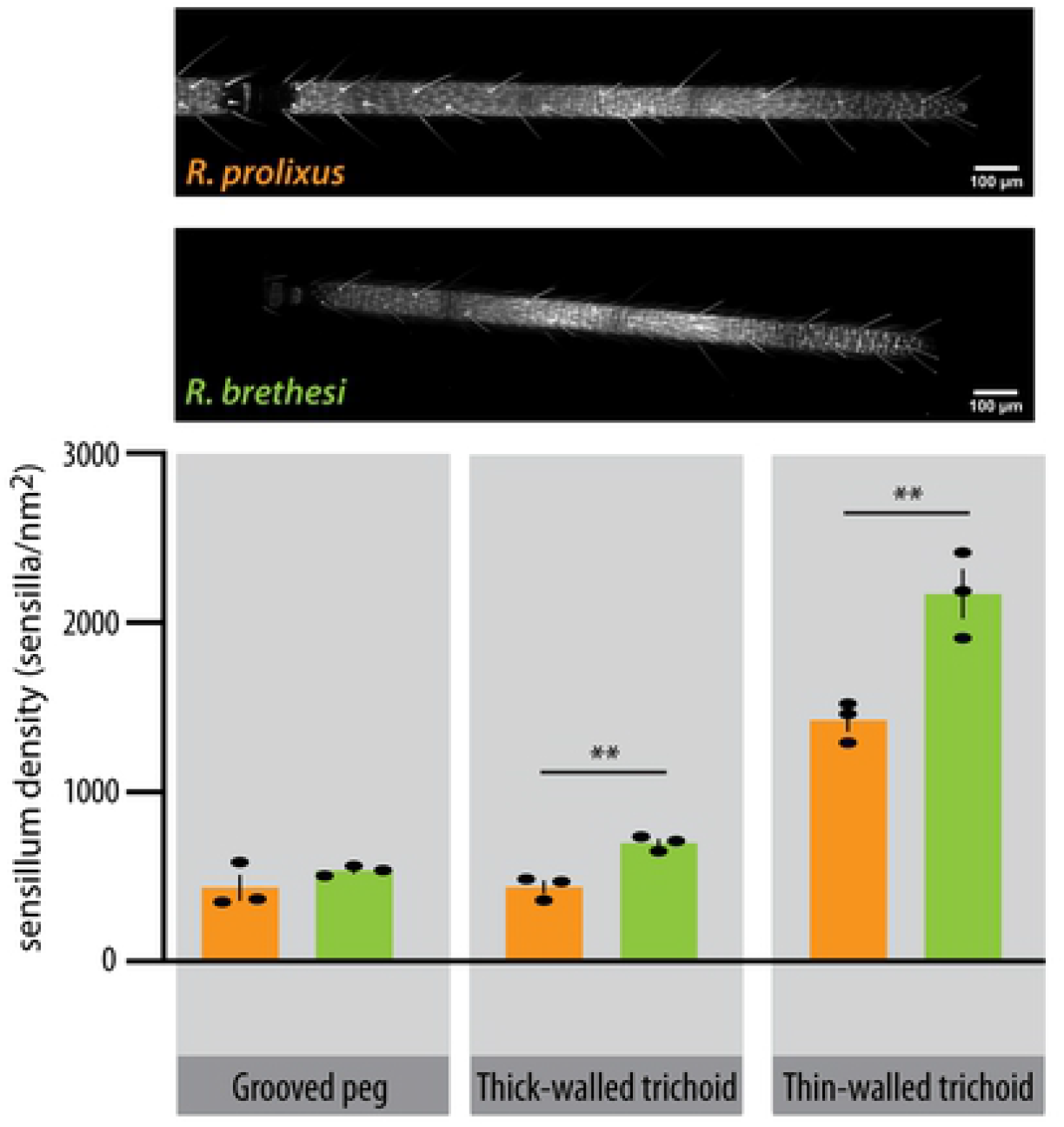
Sensillum density in domestic and sylvatic *Rhodnius* species. (A) Confocal scans from flagellomere II of *Rhodnius prolixus* and *Rhodnius brethesi* mounted in glycerol. (B) Density of olfactory sensilla in flagellomere II for each species estimated as the total number of sensilla for each type, by the flagellar surface area, n= 3, unpaired t test: **p<0.01. Data represents mean ± SEM.

### Antennal responses in Rhodnius prolixus

Previous studies on the role of olfaction in triatomines focused on *T. infestans*, by characterizing odor-evoked responses to a small number of chemical compounds, comprising acids and amines [24, 58]. We asked whether other chemical classes are also detected by the antenna of *Rhodnius* species. Thus, to identify active odor ligands we recorded, as a first approach, antennal responses using EAG recordings in *R. prolixus* to a panel of 27 odors belonging to various chemical classes and which were previously shown to elicit behavioral responses in triatomines and other haematophagous insects (**Fig. 3**, **S1 Table, S2 Table**). A significant response (i.e., p<0.05) was observed for 32% of the tested odors (one sample t test against zero). Out of these, the strongest response was observed to acetic acid, a compound that is present in triatomine feaces and mediates aggregation [59], followed by propionic acid, a known host volatile, to which *T. infestans* is behaviorally active [27]. Significant responses were also seen to the main component of the alarm pheromone [60], isobutyric acid, a compound that is also present in host volatiles [21], and to the closely related compound butyric acid. Taken together, the responses to acids represent 44% of the total significant responses. Additionally, *R. prolixus* showed a significant, though smaller, response to other host volatiles, such as cyclohexanone, amyl acetate and trimethyl amine.

**Figure 3.**
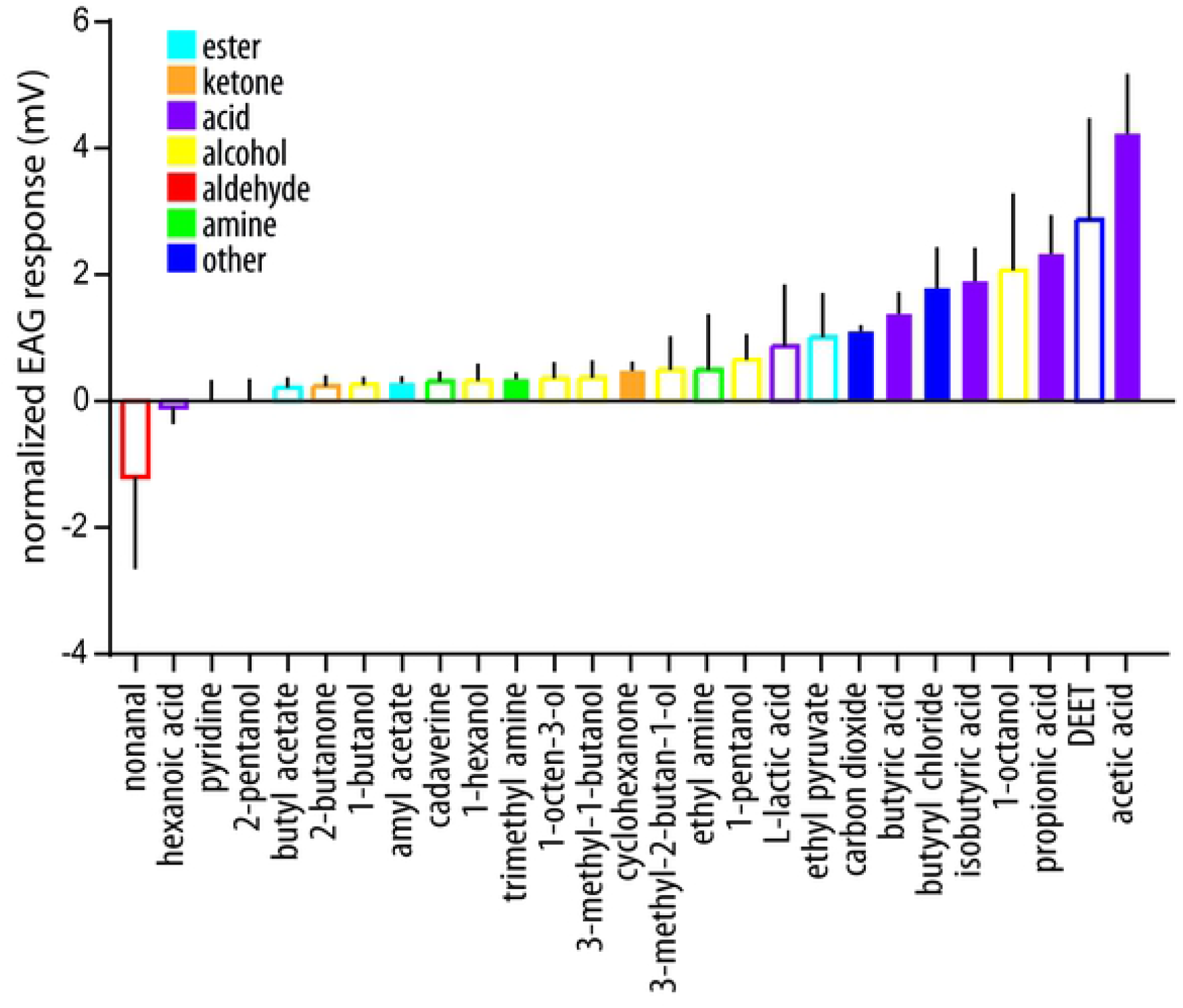
Antennal responses from *Rhodnius prolixus*. Ecologically relevant odorants (see S1 Table) were applied to the insect antenna and evoked responses were measured through electroantennogram analysis. Responses were normalized to their corresponding solvent response (see Methods). Filled bars represent statistically significant responses (one-sample t test against zero: p<0.05, n=7-10 for each odor tested).

**Figure 4.**
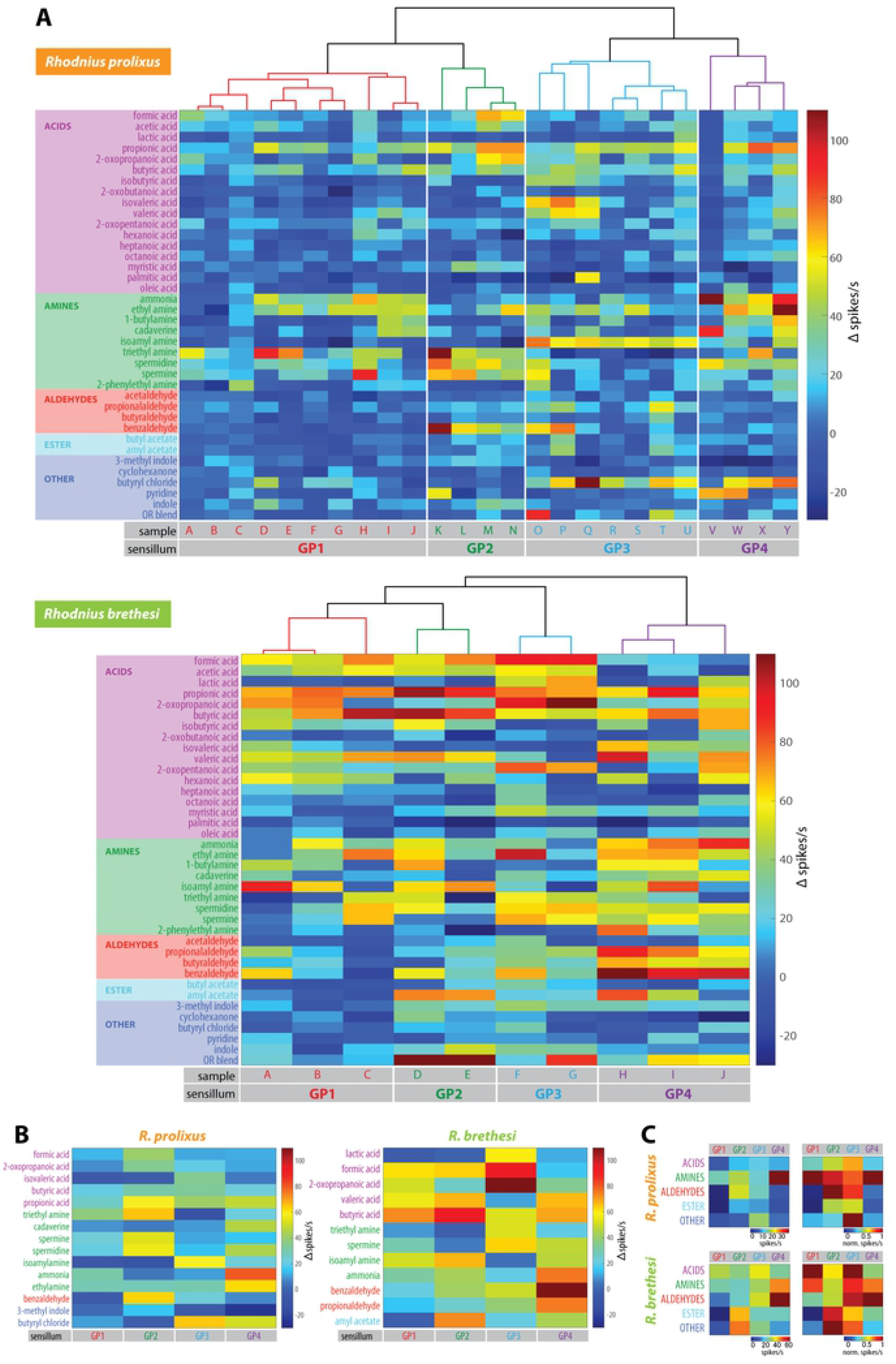
Grooved peg sensillum responses to ecologically relevant volatiles in domestic and sylvatic *Rhodnius* species. (A) Color coded responses from the grooved peg sensillum (flagellomere II) in *Rhodnius prolixus* (n=25) and *Rhodnius brethesi* (n=10). Dendrogram represents the agglomerative hierarchical clustering (Ward’s method, Euclidean distance) of these responses. (B) Color coded mean responses for the selected diagnostic odors for each sensillum type, for each insect species. Odors were chosen as diagnostic when they allowed the maximum separation between sensillum types. (C) Averaged responses for each chemical class and sensillum type (left panel), normalized to the maxiumum response within each sensillum type (right panel).

Butyryl chloride has been previously proposed as an insect repellant, as it inhibits the activity of the carbon dioxide-detecting sensory neurons in mosquitoes [44]. However, its function and detection in triatomines has not been studied so far. Interestingly, we observed a significant olfactory response to this odor (**Fig. 3**).

### Odor responses in grooved peg sensilla

Our EAG recordings of *R. prolixus* show that the olfactory system of these insects responds mostly to acids and amines. These compounds are commonly found in the environment of the insects (see **S4 Table**) and their odor-guided behavior was assessed in some species of triatomines [14, 16, 17, 39]. We next wondered whether sylvatic and domestic species present differences in their responses to these compounds. Within the antenna, acids and amines are detected by neurons housed in the grooved peg (GP) sensillum [61–63]. As shown, this sensillum type is present in the antenna of both domestic and sylvatic species of triatomines at a low density, making it an ideal system to assess differences in the olfactory tuning of insects with different habitat needs.

Each GP sensillum was screened with a comprehensive odor panel composed of compounds known to elicit a behavioral response in Triatomines and other blood-feeding insects (**S3 Table, S4 Table**). We tested a total of 38 odors, out of which 17 were acids and 9 were amines, varying in their carbon length and branches. We included additional volatiles (such as indole and amyl acetate), known to be present in, but not exclusive to, vertebrate hosts, or whose detection by the GP sensilla was shown previously in other species [19, 58]. In all, 950 odor-sensillum combinations in *R. prolixus*, and 380 odor-sensillum combinations in *R. brethesi* were tested (**Fig. 4**). We were unable to identify the number of OSNs housed in each sensillum unambiguously, therefore we defined each sensillum as a responding unit, as it has been done in studies on other insects [44]. While only 37% of the odor-sensillum combinations in *R. prolixus* yielded responses >15 spikes s^−1^ above solvent response, all combinations did in *R. brethesi* (**Fig. 4, Fig. 5**). This difference was also consistent at stronger responses (>50 spikes/s): in *R. prolixus*, only 8% of these combinations held responses higher than 50 spikes s^−1^ above solvent. In contrast, 26% of the odor-sensillum combinations in *R. brethesi* resulted in responses of >50 spikes s^−1^ (**Fig. 5A**). Notably, in both species, the odor that generated the highest number of spikes (>50 spikes s^−1^) was propionic acid (**Fig. 6C**). Responses above 100 spikes s^−1^ were generally scarce in both species.

**Figure 5.**
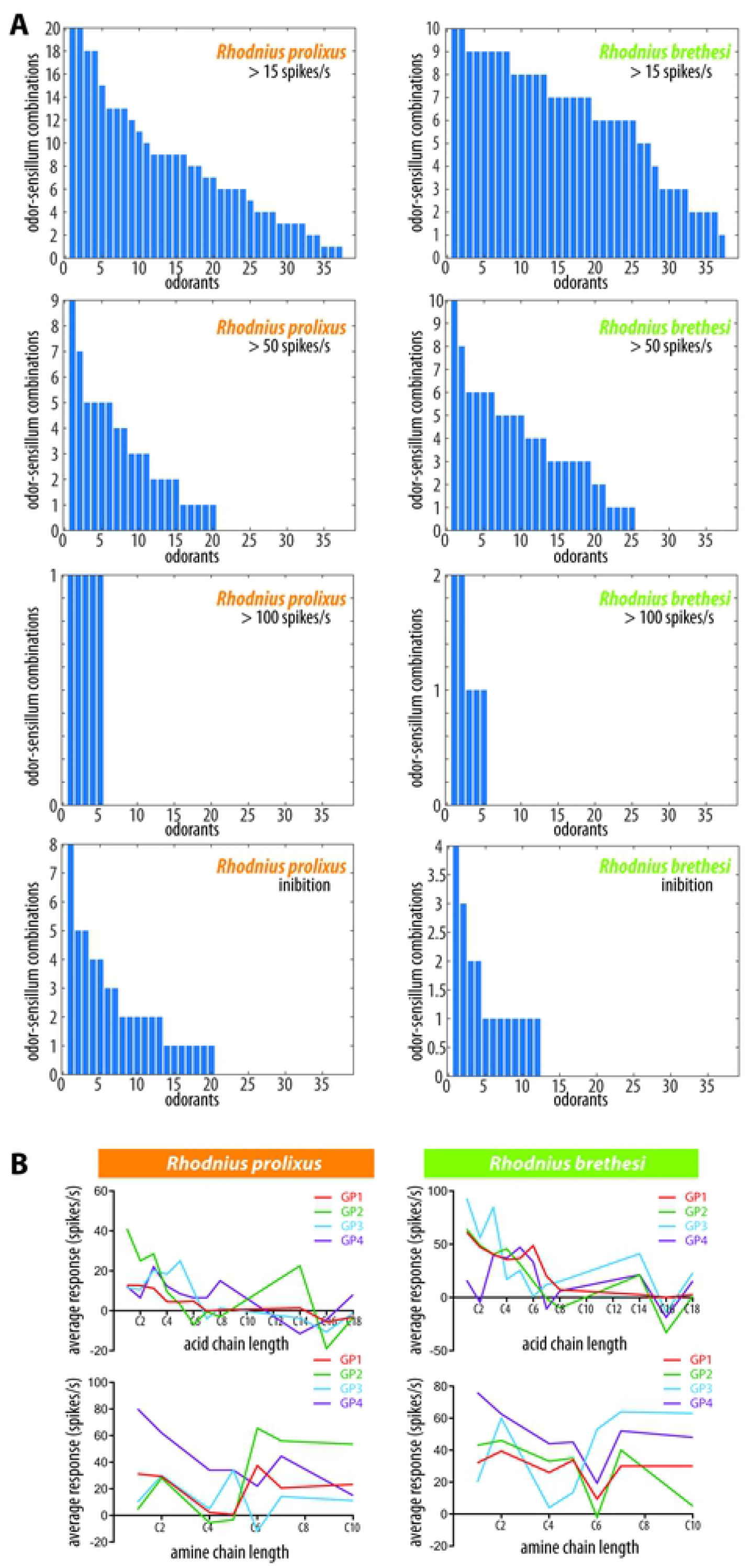
Response profile of the grooved peg sensillum. (A) Number of odor-sensillum combinations activated at the indicated firing rate by each odorant. Odorants are sorted along the x-axis according to the number of sensilla that they activate. Responses with a frequency under −15 spikes/s are considered inhibitory. (B) Averaged responses (n=2-6) to acids and amines with the indicated carbon chain length. In *R. prolixus*, GP1, GP3, GP4 show a negative correlation between carbon length and response strength to acids (Pearson correlation; Rp-GP1: r=−0.87, p=0.0005; Rp-GP2: r=−0.59, p=0.054; Rp-GP3: r=−0.75, p=0.008; Rp-GP4: −0.61, p=0.045, n=11). In *R. brethesi* GP1 and GP2 show a significant negative correlation between carbon length and response strength to acids (Pearson correlation; Rb-GP1: r=−0.89, p=0.0002; Rb-GP2: r=−0.76, p=0.006, n=11).

**Figure 6.**
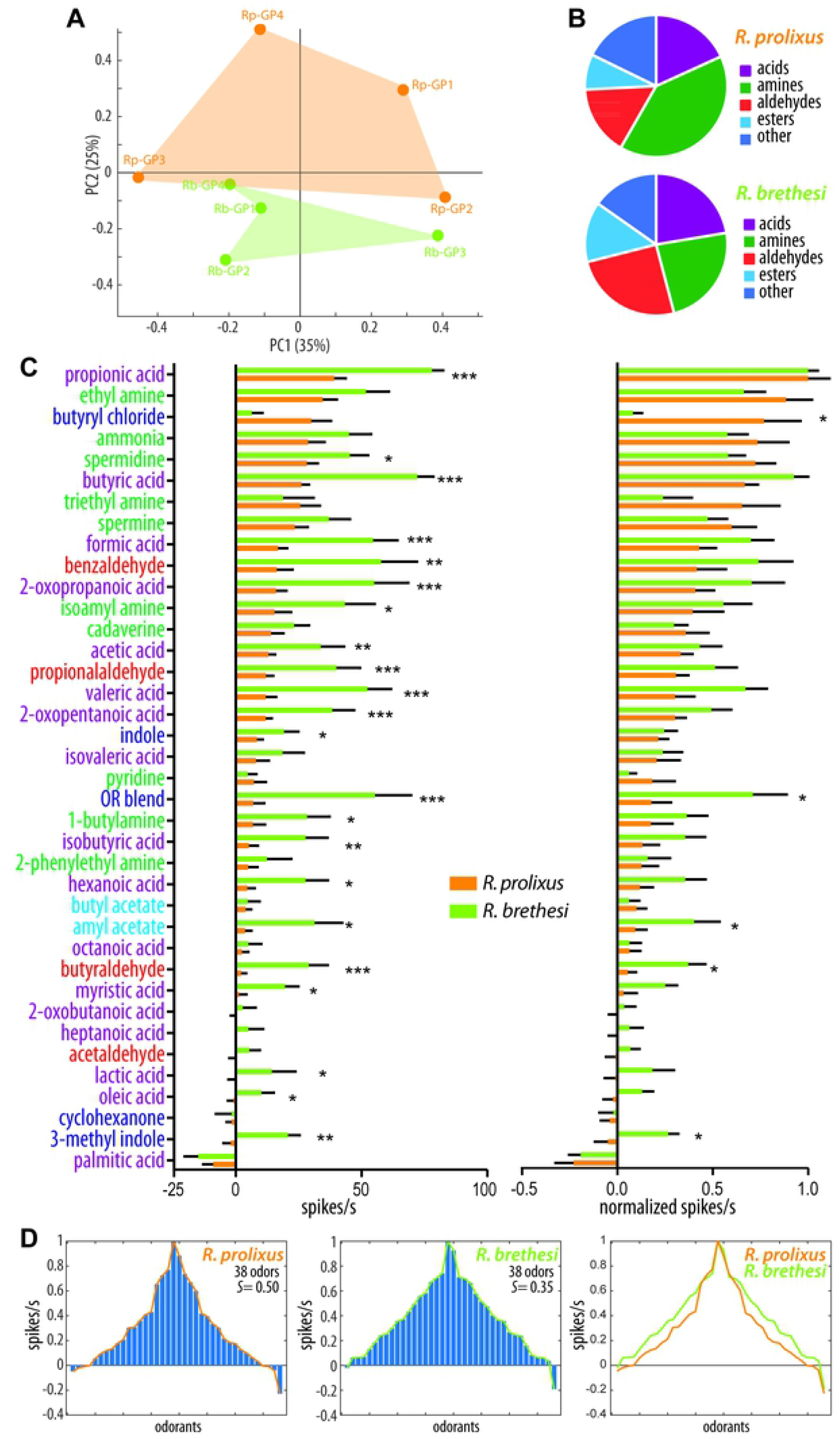
Species-specific differences: domestic species are narrowly tuned to odors. (A) Principal component analysis (PCA) of *R. prolixus* and *R. brethesi* sensillum types. No significant difference was observed for the sensillum space of the two species (ANOSIM, p=0.32, Euclidean). Sensillum responses were averaged for chemical classes (B) as well as for each odor (C), irrespective of the sensillum type to which they belong. Responses were normalized to the maximum response recorded, which was propionic acid for both species. (D) Tuning curves for each species. Odors that elicit the weakest responses are placed at the edges. The order of the odors is different for the different species. The lifetime sparseness (*S*) was calculated for each species (see Methods).

Inhibitory responses were less prevalent than excitatory ones as expected from SSR data obtained in other insect species [43] (**Fig. 5A, 6C**). In *R. prolixus*, 6%, and in *R. brethesi*, 5%, of the odor-sensillum combinations were inhibitory (<-15 spikes s^−1^ compared to solvent control). Inhibition cannot be attributed to a single odorant since 53% of the odors in the panel generated at least one odor-sensillum inhibition in *R. prolixus*, and 32% of the odors resulted in an inhibition in *R. brethesi*. However, the application of palmitic acid generated the highest number and strongest inhibitory responses in both species (**Fig. 6C**).

A major difference between the species was the response to our custom OR blend (**S3 Table**), composed of compounds typically detected by odorant receptors (ORs) in other species [40–43]. While only one sensillum responded in *R. prolixus* (> 50 spikes s^−1^) to the OR blend, 50% of the sensilla showed a response to the same blend in *R. brethesi*. Overall these results show that domestic and sylvatic triatomine species differ in their responses to the odor panel tested, with the latter showing stronger responses to a larger number of odorants.

### Funcional classification of grooved peg sensillum

To further assign the measured odor responses to distinct and functional GP sensillum subtypes in each of the two species (**Fig. 4A**), we performed an agglomerative hierarchical clustering analysis. Responses could be clustered into 4 groups in each species, corresponding to putative functional sensillum types. Each of these 4 types responded to a particular combination of odorants (**Fig. 4B**), which we propose as diagnostic odors for each specific type. In *R. prolixus*, GP type 1 (Rp-GP1), which accounts for 40% of the GP-sensilla recorded from, responds preferentially to the amines trimethylamine, ammonia and ethylamine, as evidenced by the average responses. Rp-GP2 comprises 16% of the GP-sensilla and responds best to propionic acid, triethylamine, spermine, spermidine and benzaldehyde. Rp-GP3 shows the highest responses to isoamylamine and butyryl chloride and stands for 28% of the GP-sensilla, while Rp-GP4, representing 16% of the sensilla, responds to ammonia, ethylamine and butyryl chloride.

In *R. brethesi*, the type 1 GP-sensillum (Rb-GP1) responded preferentially to butyric acid and was inhibited by amyl acetate (**Fig. 4B**). The Rb-GP2, with a similar response profile to Rb-GP1, differed from it in the responses to amyl acetate and 2-oxopropanoic acid. It also showed a higher response to isoamyl acetate, butyric, valeric and formic acids than Rb-GP1. The Rb-GP3 type showed high responses to 2-oxopropanoic acid and formic acid. Finally, type GP4 of *R. brethesi* showed a strong response to benzaldehyde, ammonia and propionaldehyde. Rb-GP1 represented 30%, Rb-GP2 20%, Rb-GP3 20% and Rb-GP4 30% of the total number of grooved peg sensilla recorded from in this species. It should be noted that all of the sensillum types presented here respond to butyric acid, as well as to propionic acid. Although these odors induced comparably higher responses, we did not include them in the suggested diagnostic panel, as they would not allow us to discriminate between sensillum subtypes.

Next, we analyzed whether certain sensillum types respond preferentially to certain chemical classes (**Fig. 4C**). In *R. prolixus*, Rp-GP1 responded strongest to amines, Rp-GP2 to aldehydes and to a lesser extent to amines. Rp-GP3 did not respond preferentially to any chemical odor class, with most responses being to butyryl chloride, and Rp-GP4 showed the strongest responses to amines. In *R. brethesi*, all of the sensillum types responded to at least two of the chemical classes tested. While both Rb-GP1 and Rb-GP3 showed the strongest responses to acids, Rb-GP3 but not Rb-GP1 responded additionally to aldehydes. Rb-GP2 did not respond to any particular odor class, with its highest responses shown to the OR blend. Finally, Rb-GP4 responded mainly to aldehydes, followed by amines.

We next evaluated whether odor compounds with a certain carbon length evoked stronger responses in the *Rhodnius* grooved peg sensilla by focusing on C1-to-C18 of acids and amines (**Fig. 5B**). In *R. prolixus*, we observed higher responses for short chain carboxylic acids (C1-6/7), with three of the sensillum types showing a negative correlation between carbon chain length and response strength (Pearson correlation; Rp-GP1: r=−0.87, p=0.0005; Rp-GP2: r=−0.59, p=0.054; Rp-GP3: r=−0.75, p=0.008; Rp-GP4: −0.61, p=0.045, n=11). Interestingly, GP2 of *R. prolixus* showed weaker responses to short chain amines, but strong ones to those with long chains (C6-C10). Acid carbon chain length also appeared to be relevant for *R. brethesi*, where it was negatively correlated with response intensity in 2 out of the 4 sensillum types (Pearson correlation; Rb-GP1: r=−0.89, p=0.0002; Rb-GP2: r=−0.76, p=0.006, n=11). In contrast, in the case of the amines, a decrease in activity with increasing carbon length was seen in GP4 (Pearson correlation; r=−0.85; p=0.016, n=7) in *R. prolixus* but not in other GP sensillum types of *R. brethesi*. However, when compared to *R. prolixus*, *R. brethesi* displayed stronger responses to short chain (C1-C5) amines (*R. prolixus:* 19.63 ± 2.65, n=125; *R. brethesi:* 19.63 ± 2.65, n=51; unpaired t test, p=0.0005).

### Species-specific differences

We next addressed the comparability of the described functional sensillum types between species. In order to get a notion of similarity between the GP types described, we calculated the Euclidean distances between the sensillum types for the two species (**S4 Table**). The averaged response values were first z-score normalized (mean=0, standard deviation=1), to ensure that the distance measured reflects dissimilarities between response patterns and not magnitude. The sensillum-pair that showed the lowest distance is GP2 in *R.prolixus* (Rp-GP2) and GP3 in *R. brethesi* (Rb-GP3; distance= 4.47). The pair Rp-GP4 and Rb-GP3 is on the other end of the spectrum, with the highest distance (8.24). In between we find most (88%) of the sensillum combinations to be within the range of 6-8.3 units of distance. In order to further explore the differences between the two species, we performed a principal component analysis (PCA) in which the 38-dimensional sensillum space was reduced to lower dimensions (**Fig. 6A**). We focused on the first two components, which together explain 60% of the variance. While the sensillum types of *R. brethesi* appeared to be more densly clustered than the ones of *R. prolixus*, the distance between individual sensillum types is larger within each species than between species (ANOSIM, R=0.09, p=0.32). This means that sensillum subtypes are not necessarily species-specific.

### Domestic species are narrowly tuned to odors

To further explore the differences between domestic and sylvatic representatives of *Rhodnius*, we averaged the responses for each chemical class (**Fig. 6B**) and odorant across sensillum types (**Fig. 6C**). While this analysis eliminates subtype specificity, it allows for a broader comparison between the species. In terms of responses to chemical classes, *R. prolixus* responded, on average, more frequently to amines than *R. brethesi*: 40% of *R. prolixus* responses were to amines, compared to 24% in *R. brethesi* (**Fig. 6B**). In contrast, *R. brethesi*, responded more strongly to aldehydes, 33% to 16%. The responses to acids within each species were comparable, with 18% of the responses in *R. prolixus* and 22% in *R. brethesi*. The same is true for the mixed chemical category (*i.e*., ‘other’): 18% of the responses in *R. prolixus* and 15% in *R. brethesi*. Finally, averaged responses of *R. brethesi* to esters were slightly higher than those of *R. prolixus*, 14% to 8%.

Averaged odor responses also stress differences between the species (**Fig. 6C**). In general, *R. brethesi* responded stronger to odors than *R. prolixus*, and a significant interspecific difference is seen for 58% of the odorants. Of those, the major differences are for the following compounds: OR blend, butyric acid, benzaldehyde, valeric acid, 2-oxopropanoic acid, propionic acid and formic acid, with *R. brethesi* presenting higher responses than *R. prolixus* in all cases. Butyryl chloride was the only compound with a significantly higher response in *R. prolixus*. When responses are normalized to the maximum average odor response (propionic acid in both species), only five of these differences remain: butyraldehyde, butyryl chloride, OR blend, amyl acetate, 3-methyl indole.

We next compared the tuning curves of the normalized odor response profiles in terms of skewness and lifetime sparseness (S) (**Fig. 6D**). This is usually done to assess how broadly or narrowly tuned olfactory receptors are. In our case this serves as a measure of the GP-sensillum tuning. As reflected in this measurement, *R. prolixus* is tuned to a narrower selection of odors than *R. brethesi*.

## Discussion

We assessed morphological and functional differences between domestic and sylvatic species of Triatominae. The antenna, the main olfactory organ, differs in the number of sensilla expressed, with a higher density of trichoid olfactory sensilla in the sylvatic *R. brethesi* compared to the domestic *R. prolixus*. Phenotypic plasticity in this character in triatomines has previously been observed [30, 31, 33, 53]. Intraspecific comparisons between insects reared under laboratory conditions and in the wild, clearly indicate that wild-caught insects often have a higher density of some types of olfactory sensilla [33, 34]. The correlation between the number of olfactory sensilla and habitat range is not exclusive to triatomines, as it has been described previously for other haematophagous insect species [52]. We observed that all but two sensillum types of *R. prolixus*, which have already been described elsewhere [24, 57, 64], were also present in *R. brethesi*. The function of two novel sensillum types is unknown, and deserves further analysis.

Using EAG recordings, we assessed the olfactory function of the *R. prolixus* antenna to known biologically active compounds, previously shown to be involved in intraspecific communication and/or in odor-guided behaviors. Surprisingly, the antenna of *R. prolixus* responded only to a limited number of the compounds tested. For instance, we did not see a significant response to either 2-butanone or 3-methyl-2-butanol, compounds known to be part of the sexual pheromone [65]. We hypothesize that the lack of antennal response is likely a consequence of the low number of specialized neurons detecting these compounds, and that the limited sensitivity of the EAG analysis failed to provide a reliable signal. However, it is conceivable that these chemicals are detected by organs other than the antenna, as the odorant co-receptor orco is expressed also in tarsi, genitalia and rosti [66]. Most of the odorants evoking a significant antennal response are volatiles characteristic of the vertebrate (amniote) odor signature, such as acetic acid, propionic acid, butyric acid, isobutyric acid, ethyl pyruvate, and trimethyl amine. All of these volatile compounds have been identified in the headspace of vertebrates, and males and females of *R. prolixus* have been demonstrated to be attracted to acetic and isobutyric acid [16]. These compounds, however, in addition to often occurring in vertebrate host secretions, are also used in intraspecific communication [67], highlighting the importance of sensory parsimony in these insects [12].

In insects, odorant (ORs) and ionotropic receptors (IRs) are responsible for the detection of volatile molecules. IRs are thought to be ancestral, as they are found in basal insects and in their most recent phylogenetic antecessor [68, 69]. These receptors, expressed in the dendrites of OSN housed in grooved peg sensilla (i.e. double walled sensilla), serve a conserved function in the detection of acids and amines across insect taxa [27, 58]. Yet, we show that triatomine insects, with different habitat and host requirements, show differences in the olfactory tuning of their GP sensilla. While both species respond to acids and amines varying in branch and carbon length, *R. prolixus* appears to be more tuned to amines than its sylvatic sibling.

In addition, domestic and sylvatic triatomine species differ in their responses to certain odorants. *R. prolixus* shows a significantly stronger response to butyryl chloride than *R. brethesi*. It is assumed that this odor compound has a repellant function in mosquitoes by inhibiting the activity of carbon dioxide-responding neurons [44]. Whether butyryl chloride serves a similar role in triatomines requires confirmation. Interestingly, *R. brethesi* revealed higher responses to amyl acetate, a compound found in fruits [70, 71], 3-methyl indole, which occurs in feces and in inflorescences at low concentrations [72–74], butyraldehyde and to the OR blend. It has been shown that compounds present in this blend are detected by odorant receptors (ORs) in other species [40–43]. While only one of the sensilla probed in *R. prolixus* responded to this blend, half of them did in *R. brethesi*. This suggests that ORs may be present in the grooved peg sensilla of *R. brethesi* but not of *R. prolixus*, as is the case, for instance, for the odorant receptor OR35a, expressed in the coeloconic sensillum of *Drosophila melanogaster* [62]. Overall, these differences might reflect specific adaptations to their corresponding environments.

Based on their odor response profiles we identified four functional sensillum subtypes in each species. This is in contrast with studies on *T. infestans*, where only three grooved peg sensillum types were described for 5^th^ instar nymphs [75]. This discrepancy might be due to several reasons. First, it is possible that differences among triatomine species are larger than expected. This is suggested by ultra-structural studies, demonstrating a different number of OSNs present in the GP sensilla of these species [45, 76]. Second, different patterns of behavior in response to odorants are also recognizable between *T. infestans* and *R. prolixus* [26]. Third, we recorded responses of adults, whereas 5^th^ instar nymphs were examined in the case of *T. infestans*. The antenna of *R. prolixus* undergoes significant changes between the 5^th^ instar and adult, with an increased number of olfactory sensilla on flagellomeres I and II [57], probably related to intraspecific communication or behavioral needs. These changes might account for the additional sensillum subtype observed in adults of *Rhodnius*. Fourth, and lastly, in our screening we recorded responses from a higher number of chemicals than in previous studies [58], potentially improving the resolution of physiological sensillum subtypes. It should be noted here, however, that our investigation is not conclusive, and recordings with additional test compounds might help to complete the ongoing work of sensilla classification in *R. prolixus* and *R. brethesi*.

Interestingly, the sylvatic species presented overall higher and broader olfactory responses than its domestic relative, as reflected in the average response, sensillum odor tuning and lifetime sparseness in SSR data. In insects, the number of olfactory receptors and the complexity of the ecological niche seem to be highly correlated, with the number of ORs increasing with niche complexity. For example, while tsetse flies have only 40-46 ORs, eusocial insects like ants possess over 350 ORs [77–79]. Notably, in triatomines, olfactory binding proteins (OBPs) and chemosensory proteins (CSPs) are present at lower expression levels in domestic insects of *T. brasiliensis*, compared to sylvatic and peridomestic ones [35]. Here we provide additional functional evidence that supports a sensory differentiation between domestic and sylvatic species. However, given the current lack of data on the chemical cues that sylvatic species encounter in the wild, we can only speculate about the selective pressures underlying these differences. It seems conceivable that the sylvatic *R. brethesi* uses olfaction to discriminate between a higher number of hosts and nest sites. In contrast, the domestic *R. prolixus* might encounter a rather limited number of olfactory stimuli. In mosquitoes, host preference has been suggested to account for differences in the chemosensory gene repertoire between sibling species [80]. Whether this is the case in triatomines we cannot confirm at present. Furthermore, studies in bat ticks suggest that host-specialization might actually be a byproduct of ecological or habitat specificity [81]. Further studies on the chemical ecology of triatomines might help to clarify this issue.

To conclude, our results confirm previous observations of phenotypic plasticity in the *Rhodnius* genus. We demonstrate that the species not only differ in the morphology of their sensory equipment, but also functionally, with domestic species presenting a distinctly decreased olfactory function, perhaps related to the limited relevance of this sensorial input in their particular environment with limited olfactory cues. It is likely that the condition found in the sylvatic species represents the ancestral character state in the subfamily, whereas a derived reduced condition is linked with a more or less close association with humans. With the ongoing rapid destruction of natural environments [8], it is likely that more species will follow this path. Careful analyses of differences and potential shifts in the sensory apparatus may turn out as helpful in the design of efficient future vector control strategies.

## Acknowledgments

We are thankful to Veit Grabe, Katharina Schneeberg, and Sandor Nietsche for help in morphological analysis, Claudio Lazzari and Marcelo Lorenzo for helpful discussions, Norisa Meli and Günther Schaub for assistance in insect breeding.

## Author Contributions

Conceived and designed the experiments: FC, SS, RI. Performed the experiments: FC. Analyzed the data: FC, SS. Contributed reagents/ materials/ analysis tools: SS, RB, BSH. Wrote the paper: FC, SS, RI, RB, BSH. Funding acquisition: FC, SS, BSH.

## Supporting information

**S1 Figure. Scanning electron microscopy (SEM) of antennal sensilla of *Rhodnius prolixus*.** Arrows indicate (A) sensillum trichobothrium (I) and bristle II (II), (B) peg-in-pit sensilla, (C) bristle III, (D) ornamented pore, and (E) type 3 coeloconic sensilla, on the pedicel of the antenna.

**S2 Figure. Scanning electron microscopy (SEM) of antennal sensilla of *Rhodnius brethesi*.** Arrows indicate (A) sensillum trichobothrium, (B) cave organ at the pedicel (I) and bristle III (II), (B’) detail of the cave organ, (C) thin-walled trichoid sensillum presenting the ecdysis channel, (D) coeloconic sensillum.

**S1 Table.** Description of the odor panel used in electrophysiological experiments.

**S2 Table.** Chemical compounds used in EAG recordings.

**S3 Table.** Chemical compounds used in SSR recordings.

**S4 Table.** Euclidean distance for the z-score normalized sensillum types of *R. prolixus* and *R. brethesi*.

